# Independent and sensitive gait parameters for objective evaluation in knee and hip osteoarthritis using wearable sensors

**DOI:** 10.1101/2020.02.20.957506

**Authors:** R.J. Boekesteijn, J.M.H. Smolders, V.J.J.F. Busch, A.C.H. Geurts, K. Smulders

**Author notes:** Address correspondence and reprint requests to: R.J. Boekesteijn, Department of Research, Sint Maartenskliniek, Hengstdal 3, 6574 NA Ubbergen, The Netherlands. Email-addresses (J.M.H. Smolders), (V.J.J.F. Busch), (A.C.H. Geurts), (K. Smulders).

## Abstract

**Objective:** To identify non-redundant gait parameters sensitive to end-stage knee and hip osteoarthritis (OA), with a specific focus on turning, dual task performance, and upper body motion in addition to straight-ahead gait.

**Design:** Gait was compared between individuals with unilateral, end-stage knee (n=25) or hip OA (n=26) scheduled for joint replacement, and healthy controls (n=27). For 2 minutes, subjects walked back-and-forth along a 6 meter trajectory making 180° turns, with and without a secondary cognitive task. Gait parameters were collected using 4 inertial measurement units on the feet and trunk. The dataset was reduced using factor analysis. One gait parameter from each factor was selected based on factor loading and effect size of the comparison between OA groups and healthy controls.

**Results:** Four independent domains of gait were obtained: speed-spatial, speed-temporal, dual task cost, and upper body motion. Turning parameters did not constitute a separate domain. From these domains, stride length (speed-spatial) and cadence (speed-temporal) had the strongest factor loadings and effect sizes for both knee and hip OA, and lumbar sagittal range of motion (upper body motion) for hip OA only.

**Conclusions:** Stride length, cadence, and lumbar sagittal range of motion were non-redundant and sensitive parameters, representing gait adaptations in individuals with knee or hip OA. Turning or dual task parameters had no additional value for evaluating gait in knee and hip OA. These findings hold promise for the objective evaluation of gait in the clinic. Future steps should include testing of responsiveness to interventions aiming to improve mobility.

## Introduction

It is well-recognized that osteoarthritis (OA) of the knee or hip impairs gait ^1–4^. Indeed, individuals with knee or hip OA walk less during daily life and their quality of gait is compromised. Yet, objective gait assessments are not part of routine clinical evaluation, and gait difficulties in OA are insufficiently captured by patient-reported outcome measures (PROMs) ^5–7^. In part, this may be due to limited time available during clinical visits, considering that gait analysis is time consuming and traditionally conducted in a gait laboratory, which is not easily accessible. Recent advances in inertial sensor technology have opened up new avenues to objectively assess gait quality in a clinical setting.

Small inertial measurement units (IMUs) can be used to quickly and accurately obtain gait parameters without being restricted to a fixed (laboratory) environment ^8^. Moreover, compared to gait analysis in a lab, substantially more strides can be collected in a shorter period of time. On the downside, an important issue of gait assessment with IMUs is that it typically results in large datasets with numerous correlated parameters, complicating the interpretation of the data. Hence, for clinical implementation, it is necessary to identify a limited set of independent gait parameters that best describe the gait adaptations in individuals with knee and hip OA compared to healthy controls.

In addition to straight-ahead level walking, turning and dual task performance have been shown to be of importance for daily life walking in elderly populations ^9–11^. These two aspects are generally overlooked in the literature regarding individuals with OA, but are meaningful to daily life ^12,13^. Turning is a common cause of falling in community dwelling elderly, and may be more sensitive to sensorimotor impairments than straight-ahead gait ^11,14^. Dual task performance, on the other hand, reflects the amount of attentional resources that are allocated to gait ^15^. In order to compensate for gait difficulties caused by OA, a strategy could be to allocate more attention to gait. The extent to which a secondary cognitive task affects gait performance (i.e. dual task cost (DTC)) may therefore be larger in individuals with OA. A third gap in literature is the lack of attention for upper body movement ^16,17^. Upper body motion is important for maintaining stability, but may also be indicative of compensatory gait changes that reflect OA-related pain or disability ^18^.

Given the low to moderate correlations between functional gait parameters and clinical or patient-reported scores, it was postulated that IMU-derived gait parameters can be supplementary in the clinical evaluation of individuals with OA ^7^. The aim of this study was therefore to identify non-redundant gait parameters that are most sensitive to the presence of knee or hip OA. In addition to spatiotemporal gait parameters, this study investigates turning, dual task performance, and upper body motion during level walking. Together, this could provide a fundament for testing responsiveness of those parameters to (non-)surgical treatment and longitudinal monitoring in OA.

## Methods

### Participants

In this case-control study, 78 subjects were included. The total study population comprised three groups: individuals with unilateral knee OA (n=25), unilateral hip OA (n=26), and healthy controls (n=27). Subjects with OA were recruited at the Sint Maartenskliniek (the Netherlands) and healthy controls were recruited from the community. Subjects were included if they had both radiological and symptomatic OA and were listed for a joint replacement surgery. Subjects had to be able to walk for more than 2 minutes without the use of any assistive device. Exclusion criteria were: 1) expectancy of joint replacement within a year, or symptomatic OA, in another weight-bearing joint than the joint scheduled for surgery, 2) BMI > 40 kg/m^2^, and 3) any other musculoskeletal or neurological impairment interfering with gait or balance. Healthy controls were recruited in the same age range as subjects with OA. Exclusion criteria for healthy controls were the same as for individuals with knee and hip OA. Informed consent was obtained from all participants prior to testing. Ethical approval was obtained from the CMO Arnhem/Nijmegen (2018-4452). All study procedures were in accordance with the Declaration of Helsinki.

### Sample size calculation

The sample size for this study was determined based on a power calculation for a longitudinal study using G*Power ^19^. This study was powered for the comparison of spatiotemporal gait characteristics between individuals 1 year after total knee or hip replacement and healthy controls. According to Senden *et al.*, effect sizes for individuals with knee OA were 1.4 for gait speed and 1.1 for stride length for this comparison ^20^. In order to achieve a power of 80%, accepting significance at 95%, with a minimum effect size of 1.1, each group required 22 subjects. Considering a 10% drop-out rate, 25 subjects in each group were required.

### Demographic and clinical assessment

Evidence for radiological OA was provided by the Kellgren and Lawrence (KL) score as assessed by experienced orthopedic surgeons (JS, VB) ^21^. Anthropometric characteristics were obtained during the pre-operative screening visit and were summarized as weight, length, and BMI. Self-reported functioning was assessed using the Knee Injury and Osteoarthritis Outcomes Score (KOOS) or Hip Disability Osteoarthritis Outcome Score (HOOS) ^22,23^. The 40 (HOOS) and 42 (KOOS) items were divided over the subscales pain, symptoms, activities of daily living, sport/recreation, and quality of life. All items were scored on a zero to four Likert scale. Total scores were transformed to a 0-100 scale, with 100 representing best function.

### Gait assessment

Gait parameters were collected on the same day as the pre-operative screening visit, which took place approximately 1 to 2 months prior to surgery. Four body-worn inertial sensors (*Opal V2, APDM Inc., Portland, OR)* were used to obtain segment accelerations and angular velocities during two gait tests (sample frequency = 128 Hz). Sensors were attached via elastic straps to the dorsum of both feet, the waist (sacrolumbar level), and the sternum (Figure 1). Subjects walked wearing their own comfortable, flat shoes (without heels). If no appropriate footwear was available, assessments were done barefoot.

**Figure 1:**
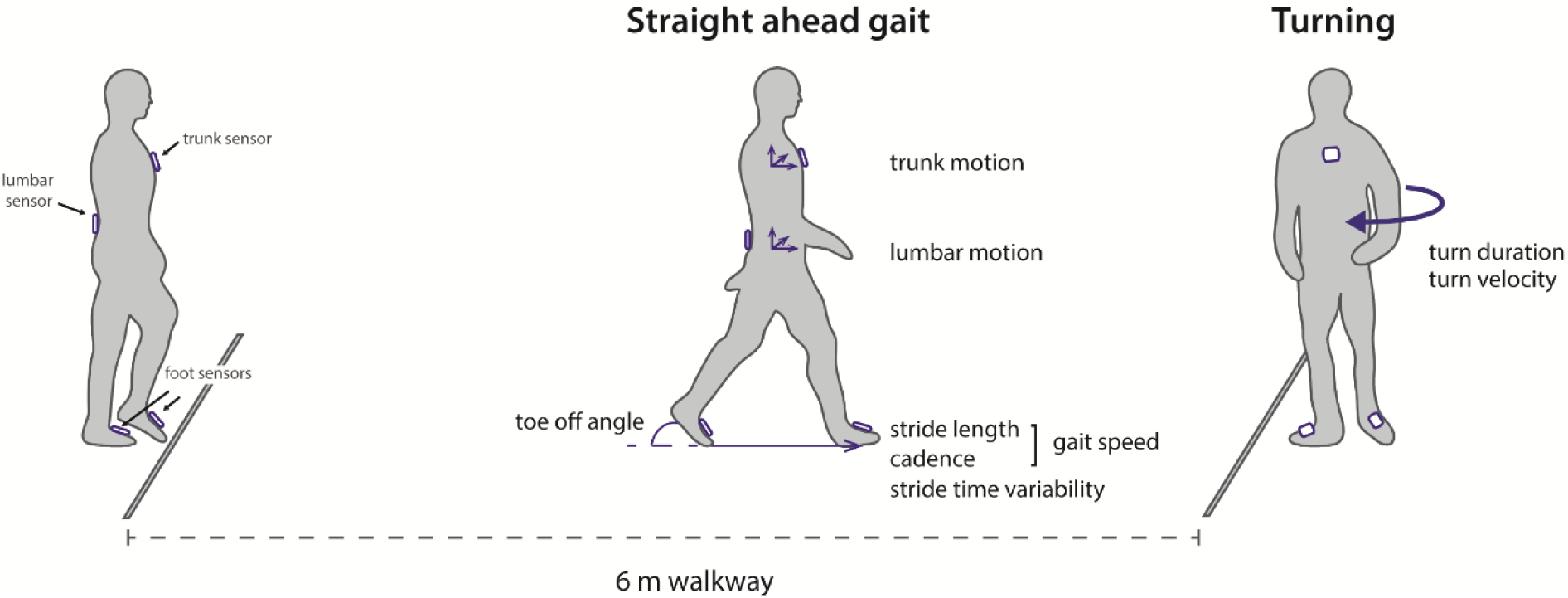
Overview of the experimental set-up. Four IMUs were attached to the dorsum of both feet, lumbar level (L4/L5) of the waist, and the sternum. For two minutes, subjects walked back and forth over the 6 meter trajectory, making 180 degree turns.

Subjects walked over a 6 meter trajectory in a corridor at the Sint Maartenskliniek. For a duration of 2 minutes, subjects walked at a self-selected comfortable speed back and forth the walkway making 180° turns (Figure 1). Two 2-minute trials were collected, with and without a secondary cognitive task. The cognitive task consisted of an alternating alphabet task, citing every other letter of the alphabet (i.e. A-C-E etc.). Single-task walking was always performed before the dual task condition. Responses to the cognitive task were recorded by the assessor. Accuracy of the alternating alphabet task during walking was summarized as the percentage of correct responses (100% *(correct responses/total responses)).

### Data analysis

Gait parameters were extracted from the raw IMU signals using the validated Mobility Lab v2.0 software package ^24^. For individuals with knee or hip OA, data from the affected leg was analyzed, whereas for healthy controls the average value from the left and right leg was taken. Gait parameters were initially selected based on their reliability ^25^, theoretical considerations, and completeness (<20% missing values). With regard to theoretical considerations, the following decisions were made: 1) in case gait parameters reflected the same outcome (e.g. gait cycle duration and cadence) only one parameter was kept for further analysis; 2) asymmetry parameters were restricted to meaningful parameters (i.e. stride length cannot be asymmetric when walking over a straight path) ^26^. DTC was computed as the percentual change of dual task performance relative to the single task for the following parameters: gait speed, cadence, stride length, stride time variability, and turn duration. DTC of gait parameters that are ratios (i.e. asymmetry values) were not included in order to prevent inflated values, except for the DTC of the coefficient of variation of stride time, due to the substantial number of other studies evaluating this parameter in the context of fall risk ^27^. This resulted in twenty-five gait parameters entered into factor analysis to identify correlated outcomes. Finally, a threshold for effect size (> 0.5) was used to obtain parameters sensitive to knee or hip OA.

Exploratory factor analysis was used to identify independent factors explaining the variance in our sample. Adequacy of the dataset for factor analysis was tested using Barlett’s test of sphericity and the Kaiser-Meyer-Olkin (KMO) test. In case individual KMO values were lower than 0.5, variables were removed from the analysis ^28^. The number of factors to be retained for further analysis was determined using the Kaiser criterium (eigenvalue > 1.0) ^29^. Subsequently, factor analysis with varimax rotation was performed to obtain orthogonal factor scores. Within a factor, gait parameters were considered relevant when they met a minimum factor loading of 0.5.

From each relevant gait parameter in the obtained factor, effect sizes were computed as standardized mean differences (SMD) for the comparison between the OA groups and healthy controls (knee OA vs healthy controls and hip OA vs healthy controls). The gait parameter with the highest factor loading in combination with an effect size larger than 0.5 for the comparison between individuals with knee or hip OA and healthy controls was considered to be non-redundant and sensitive to either knee or hip OA. For these gait parameters, individual datapoints and means with 95% confidence intervals (CI) were constructed using estimation graphs to assess between-group differences ^30^.

For demographic and clinical parameters, main group effects (3 levels: knee OA, hip OA, healthy controls) were tested using a one-way ANOVA or non-parametric equivalent when assumptions for parametric testing were not met. In cases of a significant main effect, post-hoc comparisons were conducted using independent samples Student’s t-test or the non-parametric equivalent. Data was considered statistically significant at an alpha level of 0.05. Data analysis was performed using STATA and custom-written Python scripts incorporating the DABEST library ^31^.

## Results

### Subject characteristics

Age, sex and height did not differ between OA groups and healthy controls (Table 1). Individuals with knee OA had a 9 kg (95% CI: 2 – 16; p = 0.014) higher weight compared to healthy controls. This difference was 12 kg (95% CI: 3 – 20; p = 0.007) between individuals with hip OA and healthy controls. For individuals with knee OA, this translated into a 2.8 kg/m^2^ (95% CI: 0.9 – 4.7; p = 0.005) higher BMI compared to the control group, whereas BMI was 2.4 kg/m^2^ (95% CI: 0.1 – 4.7; p = 0.043) higher in individuals with hip OA. Severity of radiographic OA was moderate to severe OA (KL = 3 or 4) in both groups. Furthermore, accuracy on the secondary cognitive task was comparable between individuals with knee (mean: 84%) or hip OA (mean: 87%) OA and healthy controls (mean: 89%). KOOS and HOOS scores indicated presence of pain, disability, and limited quality of life in individuals with knee and hip OA (Table 1).

**Table 1:** Demographic and clinical characteristics of all three subject groups.

| Parameter | Controls (n=27) | Knee OA (n=25) | Hip OA (n=26) | ANOVA main group effect | Post-hoc comparisons |
| --- | --- | --- | --- | --- | --- |
| Age (y) | 66 [63 – 68] | 64 [61 – 67] | 64 [61 – 66] | F(2,75) = 0.67, p = 0.514 | - |
| Sex (M:F) | 13:14 | 12:13 | 17:9 | $\chi^2$ (2, N=78) = 2.09, p = 0.352 | - |
| Height (m) | 1.72 [1.68 – 1.75] | 1.72 [1.68 – 1.77] | 1.76 [1.73 – 1.80] | F(2,75) = 1.72, p = 0.185 | - |
| Weight (kg) | 76 [72 – 80] | 84 [79 – 90] | 88 [80 – 95] | F(2,75) = 4.51, **p = 0.014** | Knee OA vs HC: mean diff = 9 [2 – 16 ], **p = 0.014**<br>Hip OA vs HC: mean diff = 12 [3 – 20], **p = 0.007** |
| BMI (kg/m<sup>2</sup>) | 25.7 [24.6 – 26.8] | 28.5 [26.9 – 30.1] | 28.1 [26.0 – 30.1] | F(2,75) = 3.52, **p = 0.035** | Knee OA vs HC: mean diff = 2.8 [0.9 – 4.7], **p = 0.005**<br>Hip OA vs HC: mean diff = 2.4 [0.1– 4.7], **p = 0.043** |
| KL score (I:II:III:IV) | - | 0:0:8:17 | 0:0:7:19 | - | - |
| DT scores (% correct) | 89 [86 – 92] | 84 [79 – 89] | 87 [84 – 91] | F(2,75) = 1.56, p = 0.217 | - |
| **Self-reported outcomes** |  | **KOOS** | **HOOS** |  |  |
| 1) Symptoms | - | 50.9 [42.5 – 59.3] | 41.4 [33.6 – 49.2] | - | - |
| 2) Pain | - | 41.7 [33.8 – 49.5] | 39.6 [34.4 – 44.8] | - | - |
| 3) Activities of daily life | - | 52.9 [44.9 – 60.9] | 39.7 [33.7 – 45.6] | - | - |
| 4) Sport/ Recreation | - | 15.6 [7.9 – 23.3] | 15.1 [10.5 – 19.8] | - | - |
| 5) Quality of life | - | 26.0 [20.4 – 31.6] | 23.6 [17.8 – 29.3] | - | - |

Data are presented as mean [95% CI]. Significant differences are bold.
*Note:* OA = osteoarthritis, KL = Kellgren and Lawrence, BMI = body mass index, DT = dual task, HC = healthy controls, HOOS = hip disability and osteoarthritis outcome score, KOOS = knee injury and osteoarthritis outcome score.

**Table 2:**
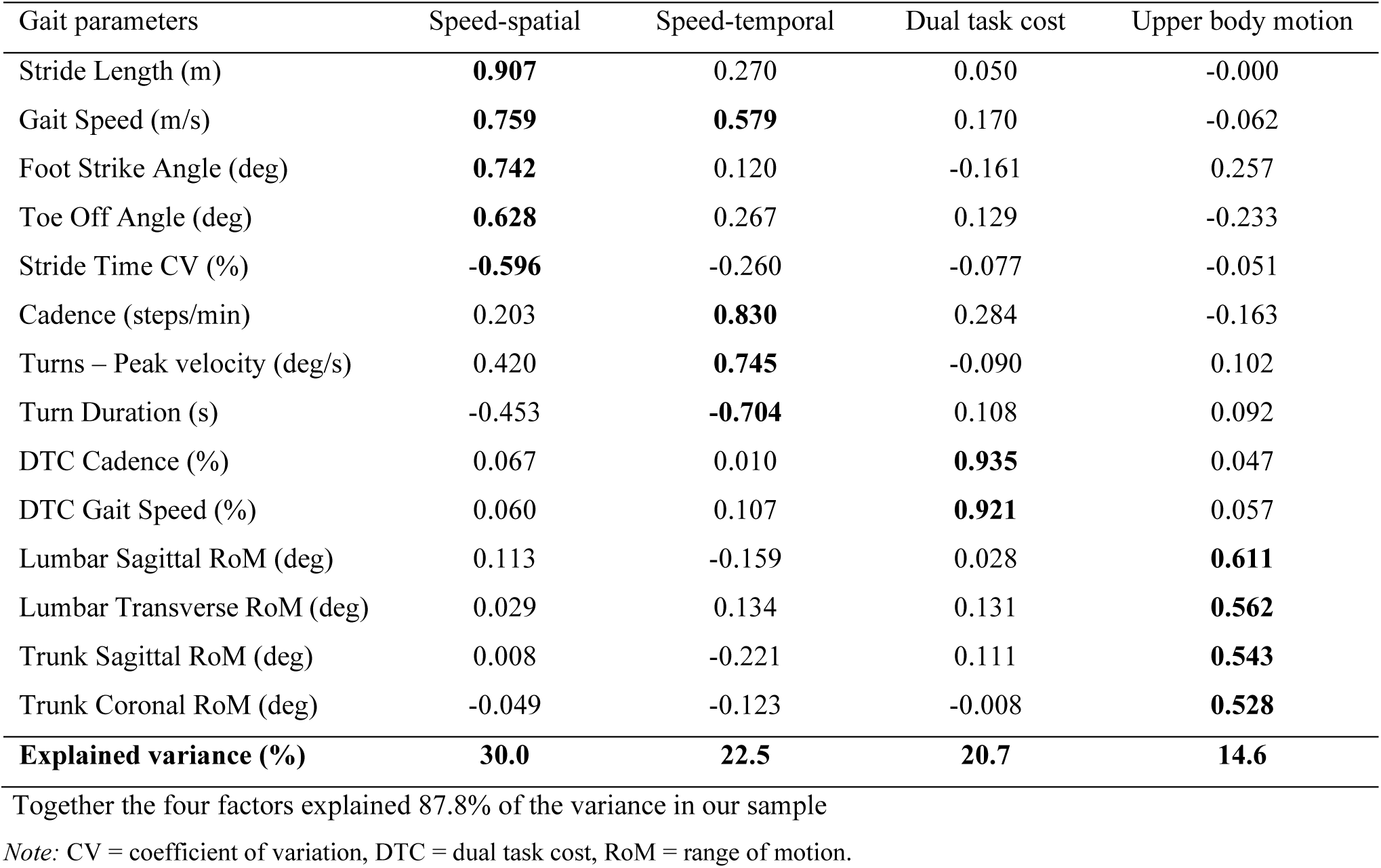
Item loadings obtained from the factor analysis (n=4) with varimax rotation.

### Exploratory factor analysis

Twenty-five gait parameters were entered into the factor analysis (Figure 2). Based on individual KMO values (<0.5) the following variables were removed from further analysis: DTC of stride length, trunk transverse range of motion (RoM), lateral step variability (spatial deviation in lateral direction of foot compared to previous steps), toe-out angle, and foot elevation at midswing (vertical foot to floor distance). The remaining twenty parameters were entered into factor analysis. Barlett’s test of sphericity confirmed the absence of an identity matrix (χ^2^ (190) = 1447.09, p < 0.001). In addition, the Kaiser-Meyer-Olkin measure was 0.666, indicating suitability of our dataset for factor analysis. Factor analysis yielded four orthogonal factors accounting for 87.8% of the total variance (Table 2). The factors were described as speed-spatial, speed-temporal, dual task cost, and upper body motion. Gait speed had a cross-loading on the factors speed-spatial (0.907) and speed-temporal (0.579). Turning parameters loaded on the factor speed-temporal and did not comprise a separate domain.

**Figure 2:**
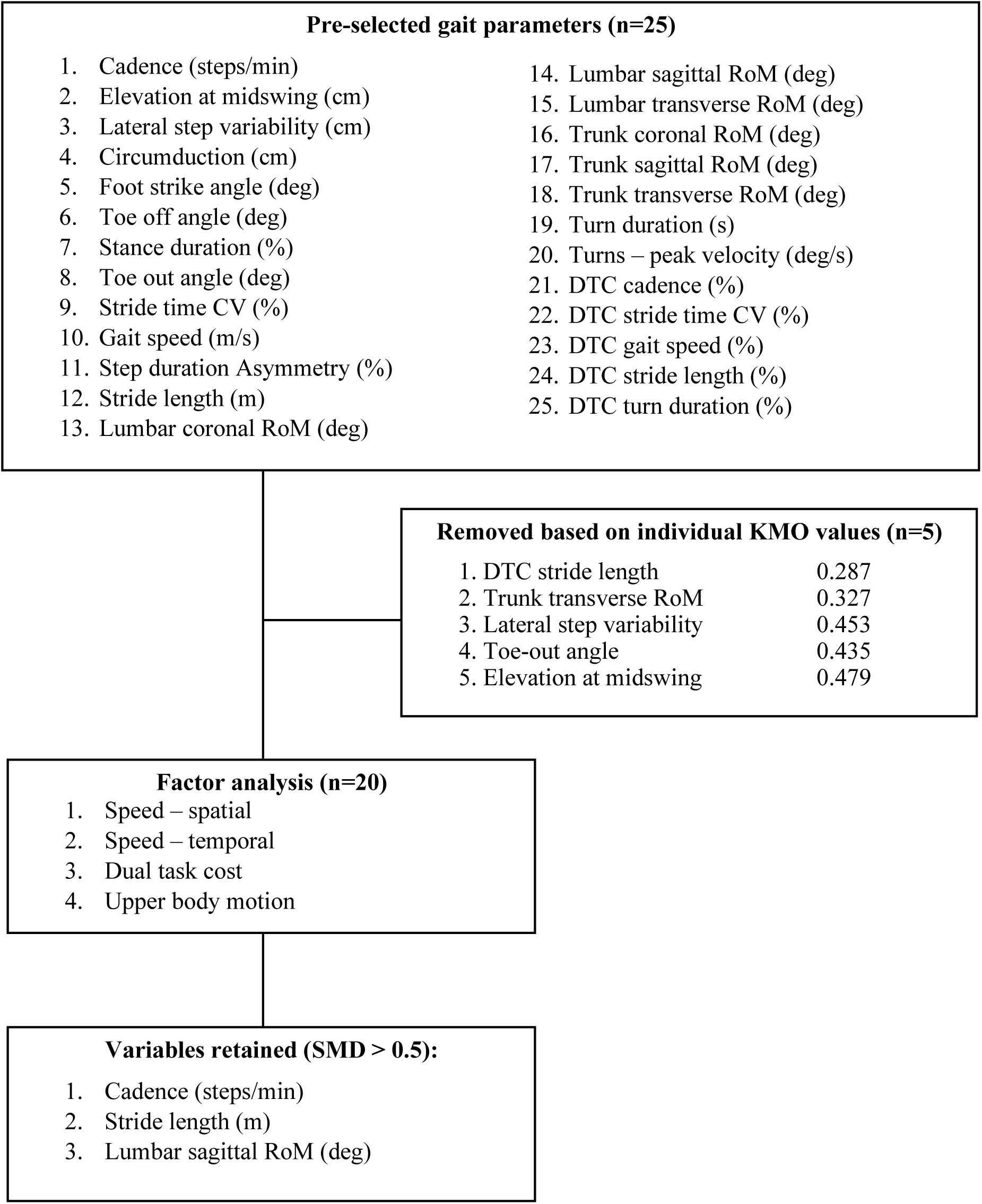
Flowchart describing the selection process of gait parameters.

### Selection of gait parameters based on effect size

Standardized mean differences for the differences between OA groups and healthy subjects for all gait parameters in the four factors are visualized in Figure 3. Based on the criterium for effect size, the following four gait parameters were selected to represent the corresponding factors: stride length (speed-spatial), cadence (speed-temporal), and lumbar sagittal RoM (upper body motion). Although the factor DTC explained 20.7 % of the total variance in our sample, none of the gait parameters within this factor showed an effect size larger than 0.5 (Figure 1). Gait speed showed the largest effect size for both the comparison between knee OA and controls (SMD = 1.59) and hip OA and controls (SMD = 1.70). However, due to cross-loadings on factors speed-spatial and speed-temporal, gait speed was not prioritized over stride length and cadence. In addition, many of the gait parameters from the factor speed-spatial and speed-temporal showed large effect sizes (SMD > 0.8) for both group comparisons.

**Figure 3:**
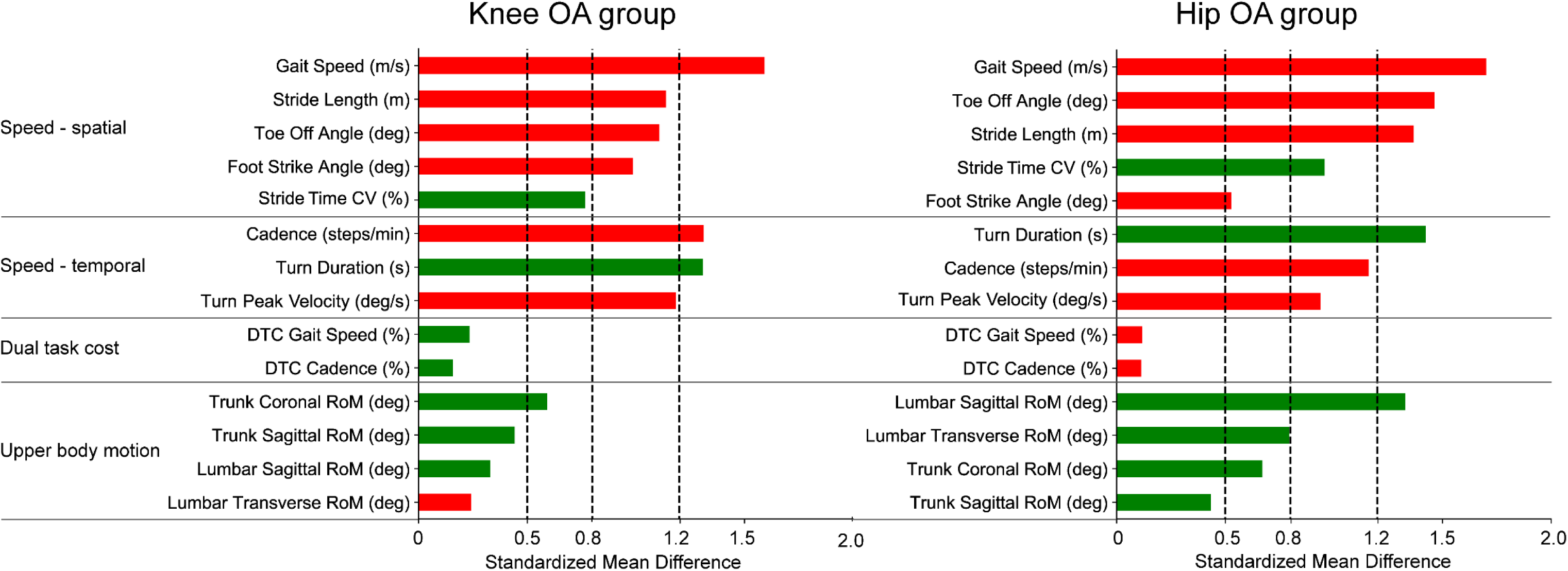
Effect sizes expressed as standardized mean differences of all gait parameters in the different factors for the comparison of healthy controls with individuals with knee OA (left) and individuals with hip OA (right). Red colors indicate OA < healthy controls, green colors represent OA > healthy controls. *Note:* CV = coefficient of variation, DTC = dual task cost, RoM = range of motion.

### Between group comparisons of non-redundant gait parameters

Between-group differences of the selected gait parameters were visualized using estimation plots (Figure 4). Both individuals with knee OA and hip OA walked with a lower cadence and with shorter steps. More specifically, compared to healthy controls stride length was 0.17 m (95% CI: 0.09-0.26, p < 0.001) lower in individuals with knee OA and 0.20 m (95% CI: 0.12-0.28, p < 0.001) lower in hip OA. In addition, cadence was 10.8 steps/min (95% CI: 6.3-15.4, p < 0.001) lower in individuals with knee OA and 9.8 steps/min (95% CI: 5.2-14.4, p < 0.001) lower in individuals with hip OA. Lumbar RoM in the sagittal plane was 2.7 degrees (95% CI: 1.7-4.4, p < 0.001) higher for individuals with hip OA compared to controls, whereas no differences were found between knee OA individuals and healthy controls (mean difference = 0.5 degrees, 95% CI: −0.33-1.59, p = 0.260).

**Figure 4:**
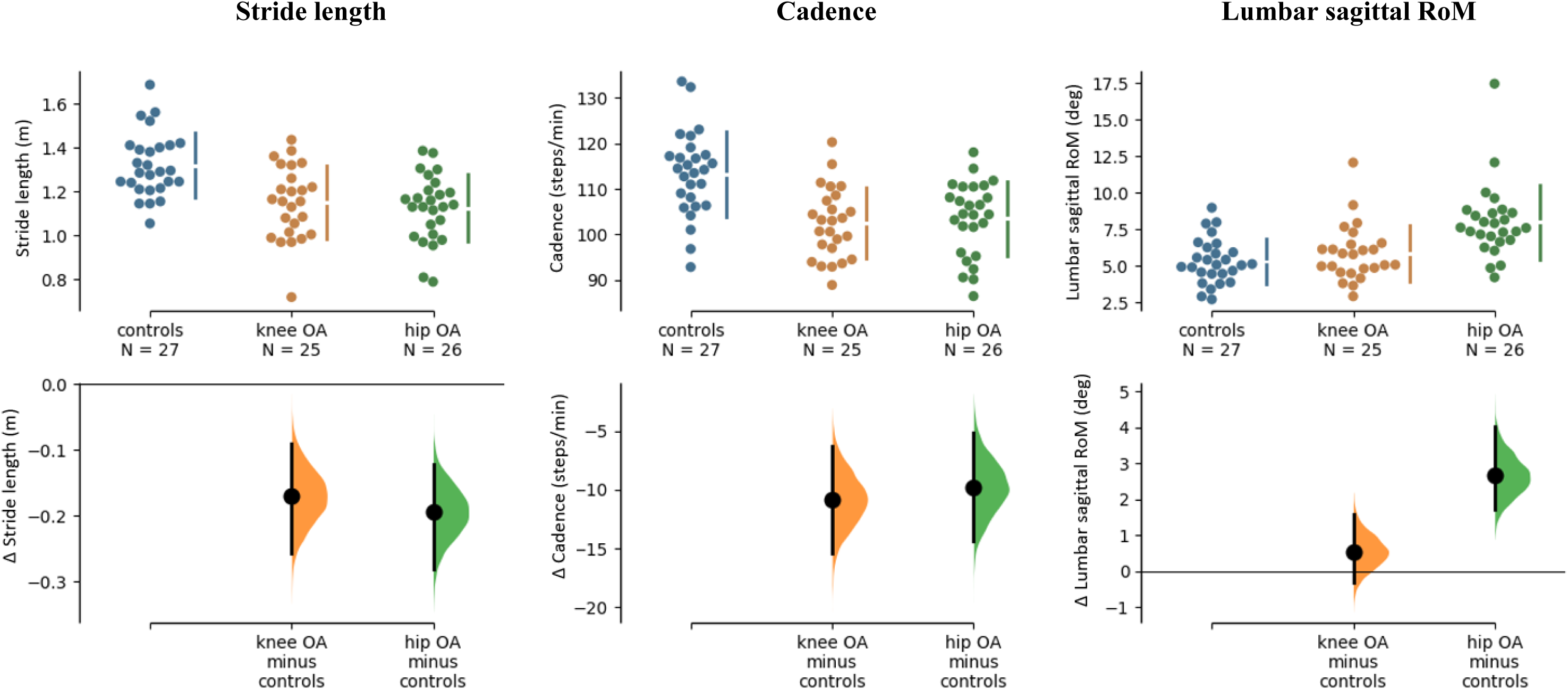
Estimation plots of the mean group differences for stride length, cadence, and lumbar sagittal RoM. In the top panel, dots represent the individual datapoints and bars the mean (± SD). In the bottom panel, the distribution of the mean difference (± 95% CI) for the comparison with healthy controls is visualized. In cases where zero is not in de 95% CI of the mean difference, as indicated by the black bars in the lower panels, data was statistically different at p < 0.05.

## Discussion

The aim of the present study was to identify a limited subset of IMU-derived gait parameters that best capture gait adaptations in individuals with end-stage, unilateral knee or hip OA. Three gait parameters were selected based on non-redundancy and sensitivity to knee or hip OA: stride length reflecting the speed-spatial domain, cadence reflecting the speed-temporal domain, and lumbar sagittal RoM reflecting the upper body motion domain. Turning or dual task parameters did not contain additional value to evaluate gait in knee and hip OA groups.

Factor analysis effectively reduced the dimensionality of our dataset from twenty-five gait parameters to four independent domains of gait. Factors reflecting the spatial and temporal aspects of gait speed are consistently reported in literature ^32–36^. Other factors related to gait reported in literature are variability ^32,33,35,36^, asymmetry ^33,35,36^, postural control ^33^, and trunk motion ^34^. Dual task cost has not previously been evaluated in a factor analysis approach. Discrepancies between our findings and factors reported in literature can be explained by the type and number of gait parameters that were initially entered for factor analysis. Importantly, dual task cost and upper body motion are interesting measures as they were independent of gait speed, evidenced by the absence of a cross-loading of gait speed on these domains in our study. Dual task cost and upper body motion may therefore provide promising gait parameters for clinical evaluation of gait in addition to speed-related measures. Turning parameters, however, seem to be redundant due to their high loading, and thus dependency, on the speed-temporal domain.

To facilitate interpretation of the gait assessment, we opted to select single gait parameters from the independent factor, to represent the respective factor. From the factors that we obtained, only dual tasking parameters did not discriminate between knee or hip OA and healthy controls (SMD < 0.5). Many of the gait parameters with large between-group effect sizes (Figure 1A and B) were grouped either under the speed-spatial or under the speed-temporal domain. This suggests that the two main components determining gait speed, stride length and cadence, are inherently linked with various gait adaptations prominent in individuals with knee and hip OA. This stresses the need to take gait speed differences into account when evaluating gait in OA. In addition, it underlines the importance of data reduction techniques, as statistical testing of all parameters would increase the probability of finding false positives.

That speed-related gait parameters have good discriminatory capacity in OA has been reported before. Two systematic reviews found a lower gait speed and stride length in individuals with hip and knee OA compared to healthy subjects ^1,3^. In studies employing IMUs, similar changes in step length and cadence were found ^20,37^. In absolute numbers, slight differences with our values can be discerned. Reasons for this may include the relatively short walking distance (6 meter) that was necessary in this study in order to obtain reliable turning measures, versus the longer distances (~20 m) that are commonly used. Nevertheless, our findings corroborated previous findings that cadence and stride length have good discriminatory capacity for both hip and knee OA.

In addition to spatiotemporal differences, individuals with hip OA walked with distinct upper body movements, which was most evident in the sagittal plane at the lumbar level. Altered movement patterns at the lumbar or thoracic level may point toward the use of compensatory strategies to unload the arthritic joint ^16^. More specifically, it was previously suggested that increased pelvic RoM in the sagittal plane may enable more effective anteflexion of the lower limbs and thereby, to a certain extent, preserves overall leg movement in the sagittal plane and thus stride length ^38^. In addition, anterior pelvic tilt combined with lateral trunk lean can reduce the lever arm between the hip joint center and center of mass ^37^. This mechanism is thought to reduce joint loading, decrease hip abductor demands during single support, and limit pain during walking. We observed more lumbar sagittal RoM and more RoM of the trunk in the coronal plane (SMD = 0.67) in individuals with hip OA compared to healthy controls. In combination this may suggest presence of lateral trunk lean and increased pelvic motion in these patients, in line with previous reports ^37,38^. Unfortunately, the exact reason for the use of these compensatory mechanisms remains speculative and may relate to pain, muscle weakness, or joint instability ^39^. Finally, compensatory trunk movements are not well reflected by a single gait parameter, as was illustrated by the relatively low factor loadings that were close together in the factor upper body motion. Future research should therefore investigate the importance of upper body motion in the different planes in individuals with OA, to inform us about potential mechanisms that may underlie these gait adaptations.

This study had several limitations that merit attention. First, we did not obtain factors representing gait asymmetry or variability, which may have been related to the low number of gait parameters related to those domains that were initially entered into factor analysis. We were therefore limited in our conclusions regarding the potential value of variability or asymmetry measures for clinical evaluation of individuals with knee or hip OA. Second, five gait parameters that could have contained valuable information were removed from further analysis due to sampling inadequacy (KMO value < 0.5). Larger sample sizes are therefore required to identify the potential value of these parameters. Finally, identifying individuals with isolated, unilateral knee or hip OA was important for our study purposes, but we acknowledge that the majority of the OA population have complaints in more than one joint ^40^. Nonetheless, we expect that widening inclusion criteria to include these patients will result in larger differences of OA groups compared to healthy controls.

In conclusion, this study provided three selected gait parameters from three independent domains (stride length, cadence, and lumbar sagittal RoM) that were sensitive to the presence of knee or hip OA. These parameters hold promise for clinical evaluation of gait in these patient groups. Adaptations in upper body motion were more subtle than stride length and cadence, but may carry important information about compensatory strategies that are distinctive for individuals with hip OA. Altogether, IMUs were well-suited to assess gait characteristics that are key for individuals with OA. Future steps should include evaluation of the responsiveness of these IMU-derived gait parameters to effects of interventions aiming to improve mobility, such as joint replacement surgery. Furthermore, longitudinal monitoring of individuals with knee and hip OA starting at earlier stages of the disease may inform us about the development of these gait adaptations and associated compensations over time.

## Acknowledgements

We want to thank Jule Mevis, Eelco van Leent, and Roosmarijn Brenninkmeijer for their contributions to the data collection.

## Author contributions

JS, VB, and KS designed the study and obtained funding. RB and KS collected the data. RB and KS analyzed and interpreted the data. JS and VB scored the radiological images. RB and KS prepared first draft version of the manuscript. AG and KS were responsible for supervision of the project. All authors read and approved the final version of the article.

## Data availability

Individual data for the gait parameters entered in factor analysis is available in the data supplement.

